# Evolutionary stasis of the pseudoautosomal boundary in strepsirrhine primates

**DOI:** 10.1101/445072

**Authors:** Rylan Shearn, Alison E. Wright, Sylvain Mousset, Corinne Régis, Simon Penel, Jean-François Lemaitre, Guillaume Douay, Brigitte Crouau-Roy, Emilie Lecompte, Gabriel A.B. Marais

## Abstract

Sex chromosomes are typically comprised of a non-recombining region and a recombining pseudoautosomal region. Accurately quantifying the relative size of these regions is critical for sex chromosome biology both from a functional (i.e. number of sex-linked genes) and evolutionary perspective (i.e. extent of Y degeneration and X-Y heteromorphy). The evolution of the pseudoautosomal boundary (PAB) - the limit between the recombining and the non-recombining regions of the sex chromosomes - is well documented in haplorrhines (apes and monkeys) but not in strepsirrhines (lemurs and lorises), which represent almost 30% of all primates. Here we studied the PAB of seven species representing the main strepsirrhine lineages by sequencing a male and a female genome in each species and using sex differences in coverage to identify the PAB. We found that during primate evolution, the PAB has remained unchanged in strepsirrhines whereas several recombination suppression events moved the PAB and shortened the pseudoautosomal region in haplorrhines. Strepsirrhines are well known to have much lower sexual dimorphism than haplorrhines. We suggest that mutations with antagonistic effects between males and females have driven recombination suppression and PAB evolution in haplorrhines. Our work supports the view that sexually antagonistic mutations have influenced the evolution of sex chromosomes in primates.

## Introduction

The human sex chromosomes are strongly heteromorphic as they exhibit extensive differences in size, gene number, DNA repeat abundance and heterochromatin composition (Skaletsky et al. 2003; Ross et al. 2005). The X chromosome comprises a large X-specific region recombining only in females whereas the Y comprises a male-specific region that does not recombine at all. Both sex chromosomes share two pseudoautosomal regions (PAR1 and 2) that recombine in both males and females. These sex chromosomes originated from a pair of identical autosomes approximately 150 million years ago, prior to the divergence of placentals and marsupials, with the evolution of Sry – the master male-determining gene in therian mammals - from Sox3 (Lahn and Page 1999; Skaletsky et al. 2003; Hughes and Rozen 2012). Since then, at several moments throughout evolutionary history, vast regions of the Y chromosome have stopped recombining with the X, likely through inversions on the Y (Lahn and Page 1999; Van Laere et al. 2008; Lemaitre et al. 2009; Pandey et al. 2013). These regions show different levels of X-Y divergence and are called evolutionary strata (Lahn and Page 1999). Strata 1 and 2 are shared among all therians, and stratum 3 is shared among all placentals (Lahn and Page 1999; Cortez et al. 2014). The most recent strata (4 and 5) have originated in the history of Catarrhini (Old World monkeys and apes) respectively, ~40 and ~25 Mya, and now only a very small PAR continues to recombine between X and Y in those primates (Hughes et al. 2012). In humans, PAR1 is the consequence of that process, while PAR2 is a recent addition (Skaletsky et al. 2003).

The process of recombination suppression between sex chromosomes, leading to a reduction in the size of the PAR and formation of evolutionary strata, has been documented in several animal and plant lineages (e.g Nicolas et al. 2005; Zhou et al. 2014; White et al. 2015). Why such a process occurred, however, is unclear. It has been proposed that sexually antagonistic mutations may have favoured the suppression of recombination (Bull 1983; Rice 1987; Charlesworth et al. 2005). Theoretical models suggest that if there are male-beneficial/female-detrimental mutations in the PAR, there will be selection to halt recombination, through for example an inversion, to genetically link those mutations to the Y chromosome,. Some evidence supporting this hypothesis has recently been found in guppies (Wright et al. 2017), but evidence from a wide range of groups, including primates, is lacking. Furthermore, there are alternative theories for why recombination is halted (reviewed in Charlesworth 2017; Ponnikas et al. 2018) and so the relative importance of sexual antagonism in sex chromosome evolution remains unclear.

While previous work on primate sex chromosomes has focused on Haplorrhini (apes, Old and New World monkeys), we studied representatives of the other main primate lineage, the Strepsirrhini (lemurs and lorises). In strepsirrhines, female social dominance (FSD), in which females dominate males, is widespread and likely ancestral (Kappeler and Fichtel 2015; Petty and Drea 2015). FSD is associated with increased testosterone production in females, resulting in the masculinization of females, including aspects of their social behaviour and genitalia (Kappeler and Fichtel 2015; Petty and Drea 2015). Some species also have rather egalitarian social systems (Pereira and Kappeler 1997). In addition, sexual size dimorphism is virtually absent among strepsirrhines (Kappeler and Fichtel 2015; Petty and Drea 2015). This is in sharp contrast with haplorrhines, where sexual dimorphism is much more pronounced and male-biased; a phenotype that is probably ancient in this group (e.g. Linderfors 2002; Kappeler and van Schaick 2004; Plavcan 2004). We therefore hypothesized that if male-female differentiation and sexually antagonistic mutations were associated with the degree of X-Y recombination suppression, strepsirrhines should show evidence of less recombination suppression compared to haplorrhines. However, to date, very little is known about the sex chromosomes of strepsirrhines, except that strata 4 and 5 are missing in gray mouse lemurs (Microcebus murinus, see Glaser et al. 1999) preventing previous tests of this hypothesis.

To identify the PAB of strepsirrhines, we used an approach relying on sequencing a male and a female at low/moderate depth, mapping the reads to a reference genome and computing the male:female depth ratio (Vicoso and Bachtrog 2011; Vicoso et al. 2013a; Vicoso et al. 2013b; Zhou et al. 2014). For autosomes, a M:F depth ratio of 1 is expected as males and females have the same copy number of autosomes. On the X chromosome, a ratio of 1 should indicate the PAR that is shared among sexes, a ratio of 0.5 should indicate the X-specific region as males have only one such region and females two, and the boundary between both would indicate the PAB. Using Illumina short-read sequencing technology, we sequenced a male and a female genome in seven species covering the main strepsirrhine lineages representing 65 My of evolution (Pozzi et al. 2014): four Lemuriformes (Daubentonia madagascariensis - aye-ayes, Microcebus murinus - gray mouse lemur, Eulemur rubriventer - red-bellied lemur, Prolemur simus - greater bamboo lemur) and three Lorisiformes (Otolemur garnetti - northern greater galago, Galago senegalensis - senegal bushbaby, Nyctibebus coucang - slow loris). The sequencing depth of each sample was between 11.8X and 39.1X (assuming a genome size identical to the human genome) with 78% of the samples being between 20X and 40X, i.e. moderate sequencing depth (Table S1). We then mapped the reads onto publicly available reference genomes of two strepsirrhines (using the human X to scaffold the strepsirrhine X chromosome) and computed a normalized M:F depth ratio to identify the X-specific region and the PAR on the X chromosome (see Methods).

## Results

Figure 1A-B shows the results for the gray mouse lemur. Using the human X chromosome to order the gray mouse lemur X scaffolds, we found that the scaffolds corresponding to human PAR1 and strata 4 and 5 have a M:F depth ratio around 1 (Fig. 1B), indicating that these regions have remained pseudoautosomal in gray mouse lemurs in agreement with older cytogenetic data (Glaser et al. 1999). The rest of the gray mouse lemur X is X-specific with a M:F ratio close to 0.5. However, five regions in the X-specific region show an elevated ratio. Detailed analysis of these five regions showed that they are fragments of autosomes (see Text S1). It is not clear, however, whether this comes from contamination of the assembly of the X chromosome by autosomal scaffolds or if this has resulted from fusion of autosomal DNA fragments to the PAR during evolution, which are misplaced in the current assembly of the X chromosome. With the fragmented assembly that is available our approach can only reliably identify the PAB, not the size of the PAR. If some autosomal material were translocated to the PAR, and thus enlarging it, it would not be possible to detect it with our approach. Only an improved assembly of the X chromosome in the gray mouse lemur could confirm one of these alternatives. Despite these limitations, it is nonetheless clear that the regions homologous to human PAR1 and strata 4 and 5 are still recombining in gray mouse lemur.

**Figure 1.**
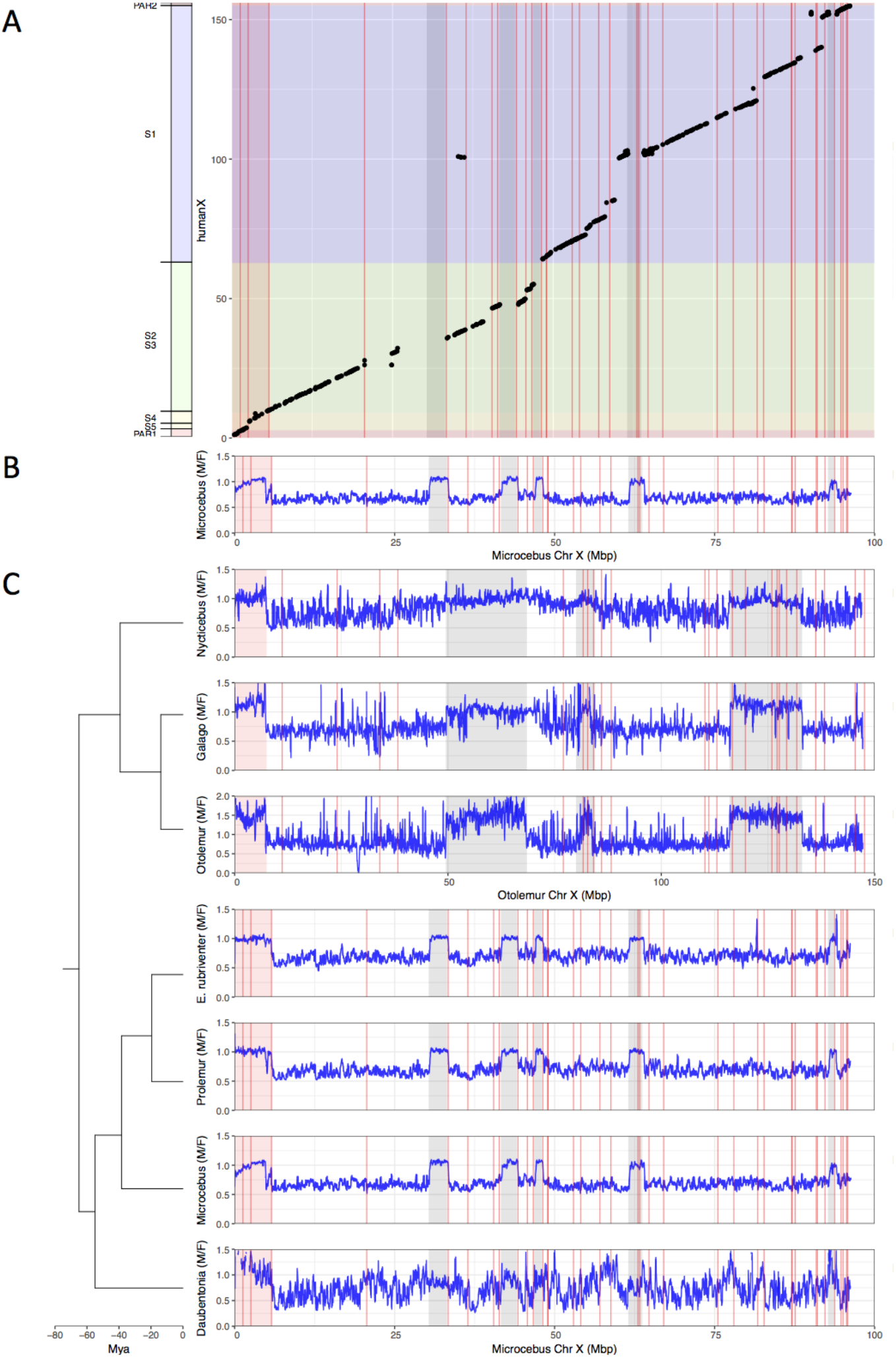
Identification the PAB in seven strepsirrhine species. (A) synteny plot of the human and gray mouse lemur X chromosomes. The human X was used to order the gray mouse lemur scaffolds (see Methods). Black dots represent orthologous genes between the human and gray mouse lemur X chromosomes. Human strata number and boundaries follow Skaletsky et al. (2003) and Hughes and Rozen (2012). Note that old strata have been split into smaller strata in Pandey et al. (2013). Human strata are indicated by different colors. S4 and S5 are in yellow. PARs are in red.(B) M:F read depth ratio along the gray mouse lemur X chromosome. Inferred PAR is shown in red. Regions of elevated M:F coverage ratio (inferred PAR plus other regions in grey) are indicated on panel A plot. (C) M:F read depth ratio for all seven strepsirrhine species. Inferred PARs for both the lemurs and the lorises are shown in red. Details on the PAR and the grey regions of the lorises can be found in Fig. S1. In all panels, red lines indicate scaffold boundaries. See Text S1 for the detailed analysis of the regions with elevated M:F coverage ratio shown in grey.

We repeated the same analysis for the other six species (Fig. 1C). For the lemurs, we used the gray mouse lemur reference genome for the mapping because it is the only one available, and for the lorises, we used the northern greater galago reference genome for the same reason (see Methods and Fig. S1 for the dot plot with the human X). Some species are quite distantly related to focal species with the reference genome and so mapping was consequently more difficult. This explains why in some cases the M:F depth ratio is more variable. The results of the aye-ayes analyses are especially noisy because of the large phylogenetic distance to the gray mouse lemur (Fig. 1C). However, in all seven species studied here, the pattern is very similar (Fig. 1C and a zoom on the PABs in Figure S2). All studied strepsirrhines harbor a large pseudoautosomal region including the genes that are in PAR1 and strata 4 and 5 in humans (compare Fig. 1A and 1C for lemurs and Fig. S1 and 1C for lorises; both lemur and loris PABs correspond to the boundary between human strata 4 and 3). We can therefore conclude that no suppression of recombination between the X and the Y has occurred in strepsirrhines since the origin of the group >65 millions years ago (Fig. 2).

**Figure 2.**
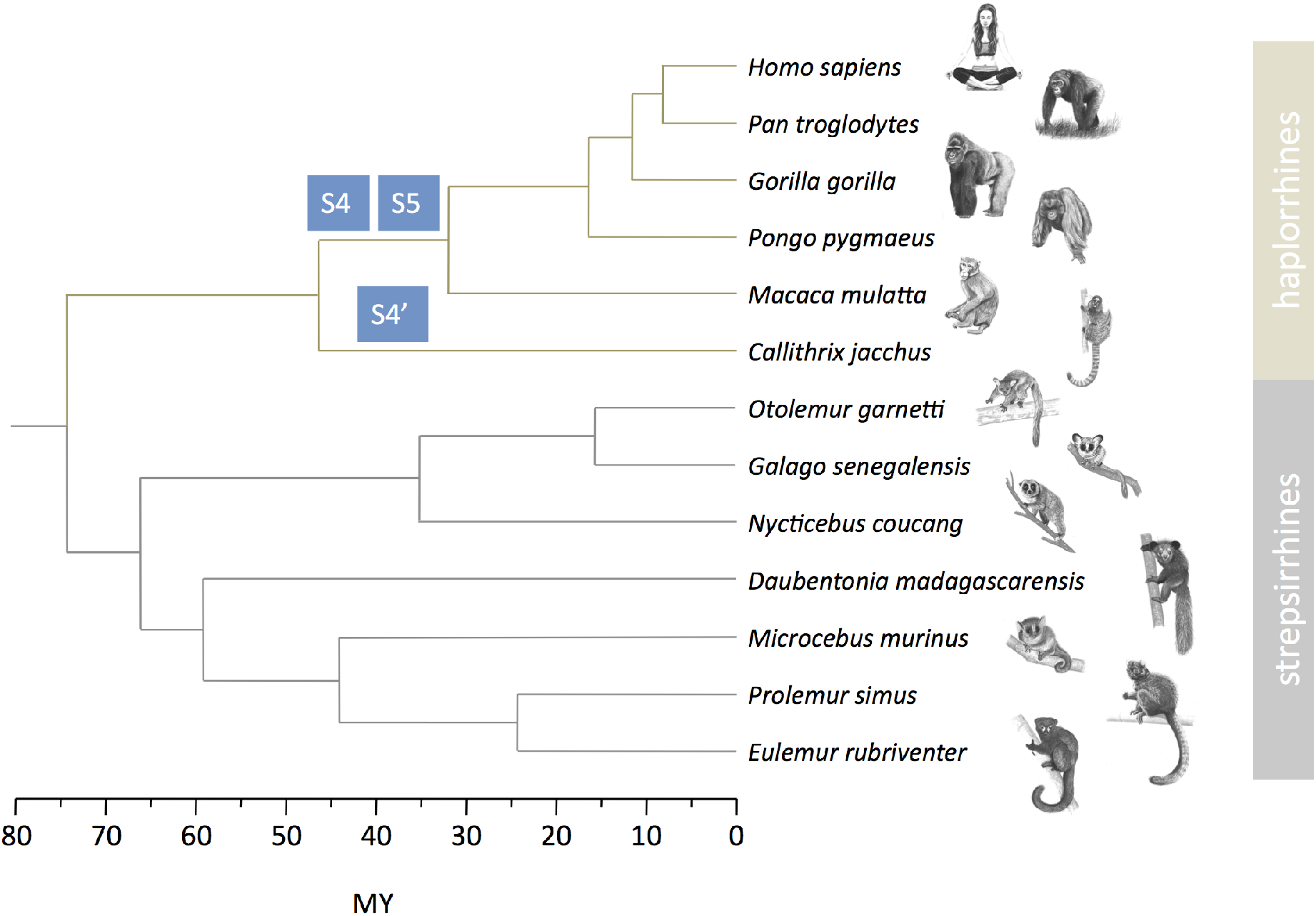
Strata formation in primates. Data on strata in haplorrhines are from Lahn and Page (1999), Skaletsky et al. (2003), Ross et al. (2005), Hughes and Rozen (2012), Hughes et al. (2012), Cortez et al. (2014). Data on strepsirrhines are from this study. The phylogenetic tree and divergence times are from Horvath et al. (2008), Pozzi et al. (2014). Drawings of primates were prepared by Philippe Faivre.

It is possible that the M:F read depth approach missed recently evolved strata in strepsirrhines. Recent strata are indeed more difficult to detect with the M:F read depth approach as sex chromosome divergence can be so low that both X and Y reads map onto the X chromosome and the ratio is close to 1 (Wright et al. 2017). To identify recent strata, we computed the male:female SNP density ratio, which is expected to more effectively detect the PAB when recent strata are present (Vicoso et al. 2013a; Wright et al. 2017). The M:F SNP density ratio is predicted to be 1 for the PAR, <1 for old strata due to haploidy in males and >1 for recent strata due to accumulation of fixed X-Y differences (Wright et al. 2017). However, our analyses revealed no recent strata in the seven strepsirrhine species studied here (Fig. S3).

Our findings are consistent with the hypothesis that recombination suppression between X and Y chromosomes is driven by sexually antagonistic mutations. However, the rate of strata formation is generally low: in primates two strata (4 and 5) were formed in apes and Old World monkeys, one was formed independently in New World monkeys (4’) based on the species studied so far (Hughes et al. 2012, Cortez et al. 2014, and see Fig. 2) and our additional data found no new strata formation in strepsirrhines. This observation is consistent with our hypothesis, but could have happened by chance because of a low common rate of strata formation in both suborders. We designed a statistical test to compare the rates of strata formation (expressed in event per My) taking into account the respective divergence times in the haplorrhine and strepsirrhine parts of the phylogenetic tree of the studied species, but this test was only marginally significant (binomial test, p=0.051 see Text S2). Because haplorrhines and strepsirrhines have different generation times, comparing rates on a generation-based timescale might however be more relevant. Rescaling time in generations to compare rates of strata formation (expressed in event per million generations) lead to a significantly higher rate in haplorrhines (binomial test, p=0.01 see Text S2), consistent with our hypothesis.

We collected phenotypic data from the literature for our set of 13 primate species and confirmed that our sets of strepsirrhine and haplorrhine species differ significantly in sexual dimorphism (teeth and body size, assuming that they reflect the global level of sexual dimorphism in an organism; see Methods and Table 1) but not in sperm competition (testes size, see Methods and Table 1).

**Table 1.** Measures of sexual dimorphism and other features in the set of studied haplorhine and strepsirrhine species.

| Species | Male Canine height in mm (sexual selection) | Female Canine height in mm (sexual selection) | Refs* | Combined Testes Mass in g (sperm competition) | Refs* | Male Body Mass in g (sperm competition) | Refs* | Male Body Mass in g (sexual selection) | Female Body Mass in g (sexual selection) | Refs* | Social and Mating System | Refs* |
| --- | --- | --- | --- | --- | --- | --- | --- | --- | --- | --- | --- | --- |
| *Callithrix jacchus* | 5,08 | 4,81 | [1] | 1,3 | [3] | 320 | [1] | 317 | 324 | [1] | Multimale | [1] |
| *Daubentonia madagascariensis* | NA | NA | NA | NA | NA | NA | NA | 2621 | 2446 | [8] | Multimale | [9] |
| *Eulemur rubriventer* | 10,49 | 9,98 | [2] | 1,76 | [3] | 2512 | [1] | 1980 | 1940 | [1] | Monogamous | [1] |
| *Galago senegalensis* | 4,01 | 3,61 | [2] | 1,66 | [3] | 210 | [4] | 227 | 199 | [2] | Unimale/Polygynous | [2] |
| *Gorilla gorilla* | 30,26 | 17,4 | [2] | 29,6 | [3] | 169000 | [1] | 170400 | 71500 | [1] | Polygynous | [1] |
| *Homo sapiens* | 10,85 | 9,97 | [2] | 40,5 | [3] | 66825 | [1] | 72100 | 62100 | [1] | Monogamous/Unimale/Polygynous | [2] |
| *Macaca mulatta* | 16,97 | 8,13 | [2] | 46 | [3] | 9200 | [1] | 11000 | 8800 | [1] | Multimale | [1] |
| *Microcebus murinus* | 2,07 | 2,08 | [2] | 2,49 | [3] | 60 | [5] | 59 | 63 | [2] | Unimale/Polygynous*** | [2] |
| *Nyctibebus coucang* | 7,05 | 6,8 | [2] | 1,2 | [3] | 1058 | [6] | 679 | 626 | [2] | Unimale/Polygynous | [2] |
| *Otolemur garnetti* | 6,53 | 6,04 | [2] | 8,93 | [3] | 320 | [7] | 794 | 734 | [2] | Unimale/Polygynous | [2] |
| *Pan troglodytes* | 21,72 | 15,26 | [2] | 128,9 | [3] | 44670 | [1] | 59700 | 45800 | [1] | Multimale | [1], [10] |
| *Pongo pygmaeus* | 27 | 15,95 | [2] | 35,3 | [3] | 74640 | [1] | 78500 | 35800 | [1] | Unimale / Multimale | [2] |
| *Prolemur simus\*\** | 5,94 | 5,91 | [2] | NA | NA | NA | NA | 2532 | 2248 | [8] | Monogamous | [2] |

\*References are [1] Lupold et al. (2019), [2] Thorén et al. (2006), [3] Harcourt et al. (1995), [4] Gomendio et al. (2011), [5] Lupold (2013), [6] Anderson et al. (2004), [7] Dixon & Anderson (2004), [8] Taylor & Schwitzer (2012), [9] Mittermeier et al. (2013), [10] Soulsbury (2010). \*\*or *Haplolemur griseus* or *H. alaotrensis* \*\*\*or multimale, see Soulsbury (2010).

## Discussion

Our work shows that, during primate evolution, the PAB has remained unchanged in strepsirrhines while several X-Y recombination suppression events have shortened the PAR in haplorrhines. We interpreted this as a consequence of differences in sexual dimorphism, and therefore sexual conflict, in both groups. However, strepsirrhines and haplorrhines differ in many ways and it is of course possible that other aspect(s) of their biology drove the pattern that we found. Strata formation may be influenced for example by gene flow (Matsumoto et al. 2017) and meiotic drive (Scott and Otto 2017) as suggested recently. Previous work has shown that the genetic diversity of strepsirrhines is highly variable (e.g Perry et al. 2012). It is however unknown whether strepsirrhines and haplorrhines exhibit systematic differences in gene flow rates and meiotic drive dynamics. A limit of this work is the use of a qualitative description of sexual dimorphism and not a quantitative one, which we could have compared to the number of strata. Future work could explore strata formation in more species and gain sufficient statistical power to compare the number of strata to phenotypic data on sexual dimorphism in primates using trait-evolution phylogenetic methods, which requires large datasets.

Evidence for the sexually antagonistic mutations hypothesis has been found in other organisms. In guppies, while the Y chromosome exhibits low levels of divergence from the X (Wright et al. 2017, Bergero et al. 2019, Darolti et al. 2019), populations exhibiting stronger sexual dimorphism seem to have a larger non-recombining region (Wright et al. 2017, Wright et al. 2019, Almedia et al 2020). In the brown alga *Ectocarpus*, sexual dimorphism is extremely low and as expected sex chromosomes are homomorphic, with a small non-recombining region, despite being very old (Ahmed et al. 2014). It should be noted, however, that other forces might be driving the process of strata formation in some lineages. In ruminants, the PAR seems to have undergone a process of attrition due to accumulation of DNA repeats (Van Laere et al. 2008; Raudsepp et al. 2015). In *Microbotryum violaceum*, strata are found on the mating type chromosomes despite the fact that this species only has mating types and not sexes, such that sexual antagonism is absent (Branco et al. 2017). Thus, sexually antagonistic mutation may not be a ubiquitous explanation of strata formation in all organisms.

Although sexual dimorphism is generally low in strepsirrhines, there are some differences among species in this lineage, with the genus *Eulemur* exhibiting the most pronounced sexual dimorphism (Petty and Drea 2015). In these species, including the red-bellied lemur (*Eulemur rubriventer*), which was analysed here, males and females exhibit striking sexual dichromatism, i.e. they differ in pelage colouration (Rakotonirina et al. 2017). The red-bellied lemur did not show more evidence for recombination suppression than the other species studied here. Sexual dichromatism may rely on sexually antagonistic mutations. The antagonism might have been solved not through Y-linkage but instead through sex-biased expression for example (Ellegren and Parsch 2007; Gazda et al. 2020). Future research could focus on sex-biased expression in strepsirrhines to test these ideas.

## Methods

### Research plan

To test whether recombination suppression is less frequent on strepsirrhine sex chromosomes compared to haplorrhines, we selected strepsirrhine species that would maximise the representation of this group’s diversity, and that were also readily accessible. We then sequenced a male and female of each species and mapped the obtained male and female reads to a reference X chromosome. The male to female depth ratio was then computed along the length of the X chromosome and the PAB was identified as the boundary between zones with a ratio of one (indicative of the PAR) and zones with a ratio of 0.5 (indicative of the non-recombining region).

### Sampling

We selected seven species covering as much phylogenetic diversity of Strepsirrhini as possible (see Table S1). Both infra-orders (Lemuriformes and Lorisiformes) are equally represented. A male and a female individual were sampled for all species (except *Otolemur garnetti*, the northern greater galago, for which sequence data from a female individual were retrieved from NCBI, see Table S1). Blood samples of *Eulemur rubriventer* (red-bellied lemur) and *Prolemur simus* (greater bamboo lemur) were collected from living animals at Zoo de Lyon in EDTA blood collection tubes to avoid coagulation. Hair samples (with follicles and roots) of the female *Daubentonia madagascarensis* (aye-aye) were collected from a living animal at Zoo Frankfurt. Samples of *Microcebus murinus* belong to the Brunoy laboratory (UMR7179, France; agreement E91-114-1 from the Direction Départementale de la Protection des Populations de l’Essonne): the biopsies were obtained from muscle tissues after the animals’ natural death. Tissues samples of a male *Daubentonia madagascariensis*, and samples of *Galago senegalensis* (Senegal bushbaby), *Nycticebus coucang* (slow loris) and of a male *Otolemur garnetti* were obtained from the tissues and cryopreserved cell collection of the National Museum of Natural History (MNHN, Paris, see Table S1).

### DNA extraction and sequencing

DNA from *Eulemur rubriventer, Prolemur simus* and female *Daubentonia madagascariensis* were extracted using two different Macherey Nagel kits. Blood samples were treated with NucleoSpin Blood Quickpure kit. Hair samples were treated with NucleoSpin DNA trace kit after a mechanical crushing of hair bulbs. DNA from the tissues and cells samples (for other species) was extracted using the DNeasy Blood and Tissue kit (Qiagen) following the manufacturer’s instructions. DNA was stored at −20° C and sent on dry ice to the sequencing platform.

A genomic DNA library was constructed for each sample using Illumina kits (TruSeq nano LT for Hiseq 2500 and 3000 sequencing). Paired-end sequencing was conducted using an Illumina Hiseq 2500 (2 x 125 bp) or 3000 (2 x 150 bp) with 1 or 2 individuals per lane at Genotoul, the INRA sequencing platform in Toulouse. Sequences were all found to be of high quality (using FastQC, https://www.bioinformatics.babraham.ac.uk/projects/fastqc) and without contamination. Consequently, no trimming was done. Sequence data and coverage are shown in Table S1.

### Chromosome assembly

Reference X chromosomes were not available for any species and genome assemblies were only available for two species that were 1) closely related to, or the same as the species being studied, and 2) assembled to an extent that it would be possible to construct a *de novo* X chromosome. These were *Microcebus murinus* (gray mouse lemur, Mmur_2.0 version from NCBI) and *Otolemur garnettii* (northern greater galago, OtoGar4 version from NCBI).

*De novo* X chromosomes were constructed for these species using scaffolds from whole genome assemblies on NCBI, which were selected, ordered and oriented against the human X chromosome. This was achieved using SynMap, an online software pipeline within the CoGe toolkit (Lyons and Freeling 2008; Lyons et al. 2008) that identified putative homologous genes between potential X scaffolds and the human X chromosome with a blast comparison (Altschul et al. 1990) using the Last algorithm (a variant of Blastz, see Schwartz et al. 2003). An algorithm within the SynMap pipeline then identified a colinear series of homologous genes between potential X scaffolds and the human X chromosome as regions of synteny, and these were arranged in order accordingly. The relative gene order DAGChainer option was used, with a maximum distance of 20 genes between two matches and a minimum of five aligned pairs of genes. The human X chromosome reference was sourced from the GRCh37.p13 Primary Assembly on NCBI (Reference Sequence: NC_000023.10).

As the results of some of the analyses in this study required normalisation using an autosome from the corresponding species, a reference autosome was constructed using the same process. In this case, the human chromosome four was used to construct a *de novo* chromosome four for *Microcebus murinus* and *Otolemur garnettii*, which was selected for its similar size to the X chromosome.

### Read mapping

Male and female reads for each species were aligned separately to their most closely related *de novo* X chromosome using Bowtie version 2-2.2.7 (Langmead et al. 2009). The reads were then sorted according to their position on the *de novo* X chromosome using Samtools version 1.3.1 (Li et al. 2009; Li 2011).

### Coverage analysis

Read depth was calculated for each sex at each position from the mapped reads on the *de novo* X using Samtools. The coverage for each sex was then normalised by dividing the depth at each position by the mean coverage depth for that species and sex on an autosome (chromosome four). The ratio of normalised male to female coverage was then calculated at each position and the data was summarised as a sliding window average using a window size of 150 kb sliding at increments of 10 kb or larger windows and increments depending on the species. This data manipulation was performed using AWK version 4.1.3.

### SNP density analysis

To detect potential regions that may have stopped recombining between strepsirrhine X and Y chromosomes relatively recently, the difference in male to female SNP density was examined for all species. For each sex of each species, SNPs were called from the mapped reads using Samtools mpileup and then converted to profiles using sam2pro version 0.8 from the mlRho package (Haubold et al. 2010). Specifically, sites with coverage <5 were excluded from the analysis and SNPs were called when a site had a minor allele frequency of 0.3 times the site coverage. The ratio of male to female SNP density was calculated for 600 kb sliding windows at increments of 10 kb. 0.001 was added to allow for a Log2 transformation and male to female SNP density was calculated at each window as Log2(sum male SNPs) – Log2(sum female SNPs). This calculation was performed using R version 3.3.2. We also calculated SNP density across an autosome (chromosome four) using the same approach and computed mean male to female SNP density and 97.5% and 2.5% quantiles across all windows.

### Statistical analysis of phenotypic differences among primates

All statistical analyses were conducted with the R statistical software (R Core Team 2019). Sexual dimorphism based on body mass (SSD, size-based sexual dimorphism) or on canine length (CSD, canine height based sexual dimorphism) was quantified as the logarithm of the ratio of the male to the female values (for instance, SSD = ln(male body mass/female body mass), Plavcan, 2004). The relative testes mass (RTM) was computed as the residual of the linear regression ln(combinedtestesmass)~ln(malebodymass).

In a first approach, the phylogenetic architecture underlying the data was ignored and we simply compared the average dimorphism value between the two groups (haplorhines vs strepsirhines). In a second stage, we accounted for the underlying phylogenetic architecture using phylogenetic contrasts in a classical phylogenetic generalized least square analysis (see Symonds and Blomberg 2014). Two evolutionary models were investigated: a simple Brownian motion (BM) and the Ornstein-Uhlenbeck model (OU) that includes stabilizing selection. The results based on the latter (OU) model should however be considered cautiously as this analysis is certainly over-parameterized considering the very small sample size (between n=11 and n=13 species). Analyses accounting for phylogenetic architecture in the data used the following specialized R packages:adephylo (Jombart and Dray 2010), ape (Paradis and Schliep 2018), geiger (Harmon et al. 2008) and phytools (Revell 2012).

Sexual dimorphism based on body mass (SSD, mean ± standard error) was 0.378±0.097 in haplorhines and 0.062±0.017 in strepsirhines. This difference based on n=13 observations was statistically significant only when ignoring phylogenetic inertia (p=0.043) but no longer significant when considering phylogenetic inertia with a Brownian motion model (p=0.66). Analysis involving an OU model would lead to a significant difference between the two groups (p=0.043) but this analysis may either be over-parameterized or suffer from the lack of phylogenetic signal in our data as revealed by the low Pagel’s λ<0.001 (not significantly different from 0) estimated in the Brownian motion model. In such a case, non phylogenetically-corrected analyses should be reported (Freckleton 2009).

Sexual dimorphism based on canine height (CSD) showed the same kind of pattern: the mean is 0.385±0.076 in haplorhines and 0.045±0.016 in strepsirhines. This difference based on n=12 observations is only significant when ignoring the underlying phylogeny (p=0.013) but no longer significant (p=0.39) when phylogeny is accounted for with a Brownian motion model (leading to a non different from 0 estimate of Pagel’s λ). The OU model leads to a significant difference between groups (p=0.013).

Based on our n=11 observations, the average relative testes mass did not significantly differ between haplorhines (0.18±0.24) and strepsirhines (−0.21±0.32). In order to avoid using residuals of a generalized least square model, we also compared testes mass in an analysis of covariance model (see Lemaître et al. 2009, for an example) including the male body mass as a covariate using the following statistical model in R: ln(combinedtestesmass)~ln(malebodymass)+group. The results were however qualitatively unchanged (the p-value associated with the “group” factor was p=0.4).

## Data accessibility

All the data generated in this study is available at NCBI (project # PRJNA482296).

## Supplementary material

Scripts for the entire coverage analysis pipeline (suitable for compute clusters using Torque job scheduling) are available on GitHub (https://github.com/rylanshearn/sex-read-depth). A supplementary material file is available at https://www.biorxiv.org/content/10.1101/445072v6.supplementary-material.

## Acknowledgements

We thank the Muséum National d’Histoire Naturelle de Paris for access to their collections of tissues and cryopreserved cells, Fabienne Aujard (UMR7179 MECADEV – CNRS, MNHN, Brunoy, France) for providing the gray mouse lemur samples and Christina Geiger from Zoo Frankfurt for providing the female aye-ayes samples. This work was performed using the computing facilities of the CC LBBE/PRABI. We thank Bruno Spataro and Stéphone Delmotte for the CC LBBE/PRABI management. We thank Peter Kappeler, Qi Zhou, Mathieu Joron and three anonymous referees for suggestions to improve this manuscript and Alice Baniel, Lounès Chikhi, Laurent Duret and Hervé Philippe for discussions. RS, EL, BCR and GABM acknowledge the financial support by ANR (grant number ANR-12-BSV7-0002-04). Version 6 of this preprint has been peer-reviewed and recommended by Peer Community In Evolutionary Biology (https://doi.org/10.24072/pci.evolbiol.100108)

