## Supplementary material file for "Evolutionary stasis of the pseudoautosomal boundary in strepsirrhine primates"

**This PDF file includes:**

Supplementary Texts S1 to S2
Figures S1 to S3
Table S1

### Supplementary Text

Text S1: Regions of the strepsirrhine X chromosomes with unusual male:female coverage ratio

In Fig. 1, both lemur X chromosomes exhibit regions with male:female coverage ratio close to 1 (shown in grey) in their X-specific parts, where a ratio of 0.5 is expected. The gray mouse lemur has five such regions, the northern greater galago three. The dot plots of the lemur and the human X chromosomes (see Fig. 1 and S1) clearly show that little or no homologous genes are found in those regions, which suggest that they may be homologous to other human chromosomes. This would be consistent with the male:female coverage ratio of 1, typical of autosomal regions, that we found for these regions. To explore this possibility, we extracted the sequences of those regions and performed a tblastn against all the human proteins (human genome version GRCh38). In case of isoforms, the longest protein was kept so that a human gene was present only once. We then filtered the tblastn results by keeping only hits with >80% similarity (based on average nucleotide divergence between lemurs and humans) and e-value < 10-9. From those, we kept human proteins covered by hits to >80% using SiLix (Miele et al. 2011). Only proteins matching to no more than one region were kept. The results of the tblastn are shown in the table below.

| | *Microcebus murinus* X chromosome regions* | | | | | *Otolemur garnetti* X chromosome regions* | | |
|---|---|---|---|---|---|---|---|---|
| Human chromosomes | 30.3-33.2 | 41.6-44.1 | 46.8-48 | 61.5-63.7 | 92.7-93.7 | 49.5-68.5 | 80-84.5 | 116-133 |
| Chrom. 1 | | | **4** | **54** | | 13 | 2 | 1 |
| Chrom. 2 | | | | 2 | | 4 | 1 | |
| Chrom. 3 | | | | | | | 1 | |
| Chrom. 4 | | | | | | 2 | | |
| Chrom. 5 | | | | 2 | | 2 | 1 | |
| Chrom. 6 | | | | | | 10 | | |
| Chrom. 7 | | | | | | | | 2 |
| Chrom. 8 | | | 1 | 1 | **4** | 1 | 1 | 1 |
| Chrom. 9 | | | | | | | 1 | |
| Chrom. 10 | | | | | | | | |
| Chrom. 11 | | | | | | | | |
| Chrom. 12 | | **8** | 1 | 2 | | 3 | **15** | |
| Chrom. 13 | | | | | | 1 | 1 | **44** |
| Chrom. 14 | | | | | | 4 | 1 | 1 |
| Chrom. 15 | | | | 2 | | 2 | | |
| Chrom. 16 | | | | 1 | | 1 | | 1 |
| Chrom. 17 | | 1 | | | | 3 | 1 | |
| Chrom. 18 | | | | | | 2 | | |
| Chrom. 19 | | | | 1 | | 3 | | |
| Chrom. 20 | | | | | | **119** | | |
| Chrom. 21 | | | | | | | | |
| Chrom. 22 | | | | 1 | | 1 | | |
| Chrom. X | | | 2 | 1 | | 2 | 3 | |

*coordinates in Mb
Human chromosome with the largest number of homologs is shown in bold

For all regions except one, most homologs that we identified are from the human autosomes, which confirms our hypothesis. These homologs are mainly from one source: chromosomes 1, 8 and 12 for regions 46.8-48, 61.5-63.7, 92.7-93.7 and 41.6-44.1 of the gray mouse lemur X chromosome, and chromosomes 12, 13 and 20 for regions 80-84.5, 116-133 and 49.5-68.5 of the northern greater galago X chromosome. These results can be interpreted two ways. One possibility is that the assemblies of the lemur X chromosome wrongly include autosomal scaffolds. Another possibility is that during the evolution of strepirrhines, some autosomal fragments have been translocated to the PAR, and the assembly failed to order these fragments correctly. Our approach cannot tell apart these possibilities but in all cases, our results suggest that these regions are probably assembly errors.

Changing tblastn outputs filtering did not change qualitatively the results. With lower %identity thresholds, we detected autosomal homologs for region 30.3-33.2 (for example, with %identity > 65, we found 2 proteins from chrom. 1, 1 from chrom. 2 and 1 from chrom 19).

Text S2: Statistical tests on strata formation

*An exact binomial test*

We partition the phylogenetic tree with total branch length $\Delta t$ into two subtrees with branch lengths $\Delta t_1$ and $\Delta t_2$, $\Delta t = \Delta t_1 + \Delta t_2$. Assuming a constant rate $\lambda$ for the formation of new evolutionary strata, the number $S$ of new strata formed during a time interval $\Delta t$ is Poisson-distributed with parameter $\lambda \Delta t$

$$\mathbb{P}(S = k) = \frac{(\lambda \Delta t)^k e^{-\lambda \Delta t}}{k!}. \quad (1)$$

On the subtree $i$, during the time interval $\Delta t_i$ we observe the formation of $S_i$ new strata. We want to contrast the following two hypotheses:

- $H_0$: Strata accumulated at a common rate $\lambda_0$ on both parts of the tree.
- $H_1$: Strata accumulated at different rates $\lambda_i$ during the time intervals $\Delta t_i$.

We use the number $S_1$ of strata formed in the time interval $\Delta t_1$ as the test statistics and compute the conditional probability to observe a larger value given the total number $S_1 + S_2$ of strata formed in the time interval $\Delta t_1 + \Delta t_2$ under the null hypothesis $H_0$:

$$\begin{aligned}\mathbb{P}(S_1 \geqslant k_1 | S_1 + S_2 = k_1 + k_2, \Delta t_1, \Delta t_2) &= \frac{(k_1+k_2)!}{(\lambda_0(\Delta t_1 + \Delta t_2))^{k_1+k_2} e^{-\lambda_0(\Delta t_1+\Delta t_2)}} \sum_{j=k_1}^{k_1+k_2} \frac{\lambda_0^{k_1+k_2} \Delta t_1^j \Delta t_2^{k_1+k_2-j} e^{-\lambda_0(\Delta t_1+\Delta t_2)}}{j!(k_1+k_2-j)!} \\ &= \sum_{j=k_1}^{k_1+k_2} \binom{k_1+k_2}{j} \left(\frac{\Delta t_1}{\Delta t_1 + \Delta t_2}\right)^j \left(\frac{\Delta t_2}{\Delta t_1+\Delta t_2}\right)^{k_1+k_2-j},\end{aligned} \quad (2)$$

where we recognize the binomial distribution. Note that this probability is independent of the common rate $\lambda_0$ of the Poisson process. Applying this test is conceptually equivalent to tossing an unbalanced coin $k_1 + k_2$ times with a probability $p = \frac{\Delta t_1}{\Delta t_1 + \Delta t_2}$ to get a head and computing the probability to obtain at least $k_1$ times a head.

*Evolutionary times*

The phylogenetic relationships and mean divergence times for the included primate species were recovered from the previously published primate phylogeny and divergence dates (Pozzi et al., 2014, supplementary table 3). Detailed phylogenetic relationships among strepsirrhine lineages (Horvath et al., 2008) were used to infer phylogenetic reletionships in the cases when species in our analysis were not included in this reference study. The divergence times are shown in a separate table.

Generation times in the studied primate species are highly variable, and we are mostly interested in comparing *per generation* rather than *per year* rates of strata formation. For this purpose, the branch lengths in the phylogenetic tree need to be rescaled by the generation times. We used the age at first reproduction as a proxy for generation time following the example of Gaillard et al. (2005). Ages at first reproduction for the extant species in the phylogenetic trees were obtained from Ernest (2003) and maximum likelihood estimates of this trait were obtained for internal nodes of the phylogenetic tree with the `fastAnc` method implemented in `phytools` (Revell, 2012). This method assumes that the age at first reproduction evolves neutraly according to a Brownian motion model (Felsenstein, 1973; Schluter et al., 1997).

The branch lengths of the phylogenetic tree were rescaled by the generation times. In order to take into account variable generation time along a branch, we used the following method: We denote $g$ the time counted in generations and $t$ the time counted in years along a phylogenetic branch. The instantaneous generation time (expressed in years per generation) along a given branch at any time $t$ is $\gamma(t) = \frac{dt}{dg}$. We assume a linear trend for $\gamma(t)$ between an ancestral node (at $t = t_a$, for which $\gamma(t_a) = \gamma_a$) and a descendant node (at $t = t_d$, for which $\gamma(t_d) = \gamma_d$). This assumption raises the following ordinary differential equation:

$$\gamma(t) = \frac{dt}{dg} = \gamma_a + \lambda(t - t_a),$$

where $\lambda = \frac{\gamma_d - \gamma_a}{t_d - t_a}$. The general form for the solutions of this equation is

$$g(t) = \frac{1}{\lambda} \ln(\gamma_a + \lambda(t - t_a)) + K,$$

where $K$ is an integration constant. The number of generations elapsed on the branch between times $t_a$ and $t_d$ is thus

$$g(t_d) - g(t_a) = \frac{t_d - t_a}{\gamma_d - \gamma_a} \ln \frac{\gamma_d}{\gamma_a}. \quad (3)$$

The ages at first reproduction for the extant species and their maximum likelihood estimates as well as the rescaled branch lengths in the primate phylogeny are shown in a separate table.

*Results*

The haplorrhine lineages in our sample have evolved for for $\Delta t_1 = 188.52$ My (44.23 million generations) during which $S_1 = 3$ new strata were formed. The strepsirrhine lineages evolved for $\Delta t_2 = 321.32$ My (158.52 million generations) and no new strata was formed ($S_2 = 0$). Comparing the rates of strata formation expressed in number of events per million year leads to a marginally significant $p$-value (one-tailed binomial test, $p = 0.051$), this trend becomes significant when considering the rates expressed in number of events per million generations (one-tailed binomial test, $p = 0.010$) suggesting that new strata form at a higher rate per generation in the haplorrhine lineages.

**Table: Divergence time estimates in the primate phylogeny.**

| Node label[a] | Mean age[a] (My) | Descendant nodes or species | |
|---|---|---|---|
| 4 | 7.65 | *Homo sapiens* | *Pan troglodytes* |
| 5 | 10.63 | 4 | *Gorilla gorilla* |
| 7 | 17.29 | 5 | *Pongo pygmaeus* |
| 38 | 32.12 | 7 | *Macaca mulatta* |
| 44[b] | 46.72 | 38 | *Callithrix jacchus* |
| 61 | 74.11 | 44 | 60 |
| 50[c] | 24.24 | *Eulemur rubriventer* | *Prolemur simus* |
| 54[d] | 43.46 | 50 | *Microcebus murinus* |
| 55 | 59.55 | 54 | *Daubentonia madagascarensis* |
| 56 | 17.28 | *Galago senegalensis* | *Otolemur garnetti* |
| 58 | 36.35 | 56 | *Nycticebus cougang* |
| 60 | 66.33 | 55 | 58 |

[a]For sake of clarity we report the node labels and ages from our reference primate phylogeny (Pozzi et al., 2014).

[b]*Callithrix jacchus* is not included in the reference phyolgeny but the divergence time between platyrrhini and catarrhini can be used.

[c]*Prolemur simus* and *Eulemur rubriventer* are not included in the reference phylogeny, but the divergence between *Eulemur macaco* and Lemur catta can be used (node 18 in Horvath et al., 2008).

[d]*Microcebus murinus* is not included in the reference phylogeny, but the divergence with *Lepilemur sp.* can be used (node 7 in Horvath et al., 2008).

**Table: Rescaling of the primate phylogeny with generation time.**

| Ancestral node | | Descendant node | | Branch length | |
|---|---|---|---|---|---|
| Node[a] | AFR[b] | Node[a] | AFR[b] | My | MGen |
| 4 | 119.12 | *H. sapiens* | 131.87 | 7.65 | 0.73 |
| 4 | 119.12 | *P. troglodytes* | 131.87 | 7.65 | 0.73 |
| 5 | 109.18 | 4 | 119.12 | 2.98 | 0.31 |
| 5 | 109.18 | *G. gorilla* | 89.33 | 10.63 | 1.29 |
| 7 | 99.42 | 5 | 109.18 | 6.66 | 0.77 |
| 7 | 99.42 | *P. pygmaeus* | 110.12 | 17.29 | 1.98 |
| 38 | 68.49 | 7 | 99.42 | 14.83 | 2.14 |
| 38 | 68.49 | *M. mulatta* | 41.08 | 32.12 | 7.19 |
| 44 | 50.51 | 38 | 68.49 | 14.6 | 2.97 |
| 44 | 50.51 | *C. jacchus* | 16.33 | 46.72 | 18.52 |
| 61 | 36.80 | 44 | 50.51 | 27.39 | 7.59 |
| Total length for the haplorrhine subtree ($\Delta t_1$) | | | | 188.52 | 44.23 |
| 61 | 36.80 | 60 | 32.91 | 7.78 | 2.68 |
| 60 | 32.91 | 58 | 23.70 | 29.98 | 12.82 |
| 60 | 32.91 | 55 | 31.60 | 6.78 | 2.52 |
| 58 | 23.70 | 56 | 17.99 | 19.07 | 11.05 |
| 58 | 23.70 | *N. cougang* | 23.42 | 36.35 | 18.51 |
| 56 | 17.99 | *O. garnetti* | 19.48 | 17.28 | 11.07 |
| 56 | 17.99 | *G. senegalensis* | 11.33 | 17.28 | 14.40 |
| 55 | 31.60 | *D. madagascarensis* | 39.00 | 59.55 | 20.32 |
| 55 | 31.60 | 54 | 26.49 | 16.09 | 6.67 |
| 54 | 26.49 | 50 | 26.39 | 19.22 | 8.72 |
| 54 | 26.49 | *M. murinus* | 12.90 | 43.46 | 27.61 |
| 50 | 26.39 | *P. simus* | 28.93 | 24.24 | 10.52 |
| 50 | 26.39 | *E. rubriventer* | 23.73 | 24.24 | 11.62 |
| Total length for the strepsirrhine subtree ($\Delta t_2$) | | | | 321.32 | 158.52 |

[a]For sake of clarity we report the node labels of our reference primate phylogeny (Pozzi et al., 2014).

[b]Values of the age at first eproduction (AFR, expressed in months) for extant species are from Ernest (2003). The value for *H. sapiens* was not available and conservatively set to the same value as for *P. troglodytes*. The value for *E. rubriventer* was not available and was estimated as the mean value of other species in the same genus. The values for internal nodes of the primate phylogeny are maximum-likelihood estimates under a Brownian motion model (Felsenstein, 1973; Schluter et al., 1997) obtained with `phytools` (Revell, 2012).

**Fig. S1.**

Synteny analysis of northern greater galago and human X chromosomes. (A) synteny plot of the human and northern greater galago X chromosomes. (B) M:F read depth ratio along the northern greater galago X chromosome. See legend of figure 1 for more details.

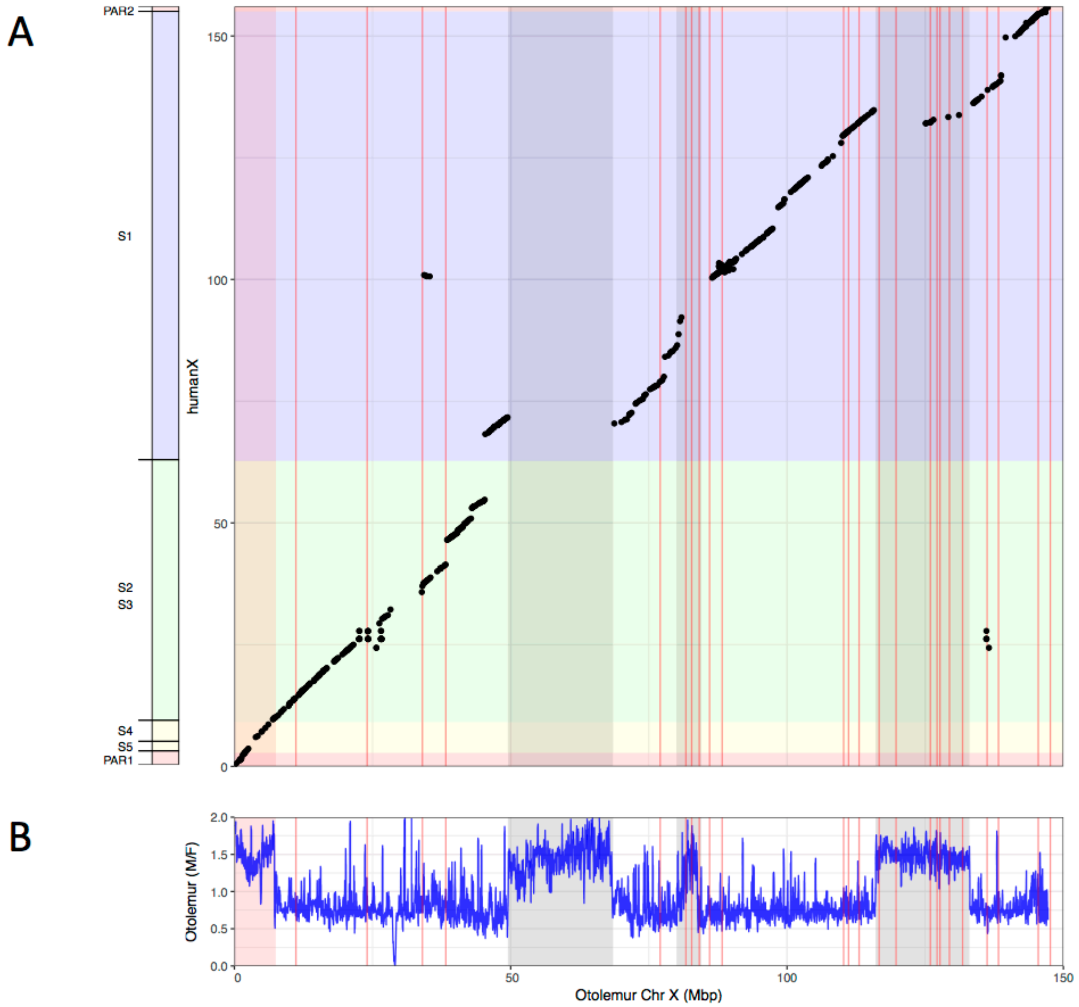

**Fig. S2.**

Zoom-in on the inferred PABs. (A) combined M:F read depth ratio for northern greater galago (red), senegal bushbaby (green), slow loris (blue). (B) combined M:F M:F read depth ratio for aye-ayes (red), gray mouse lemur (blue), red-bellied lemur (light blue), greater bamboo lemur (purple). Position of the PABs in lemurs and lorises is the same (see legend of figure 1 for more details). Positions of the PABs in Mb shown here differ because of differences in the X chromosome assembly between *M. murinus* and *O. garnetti*.

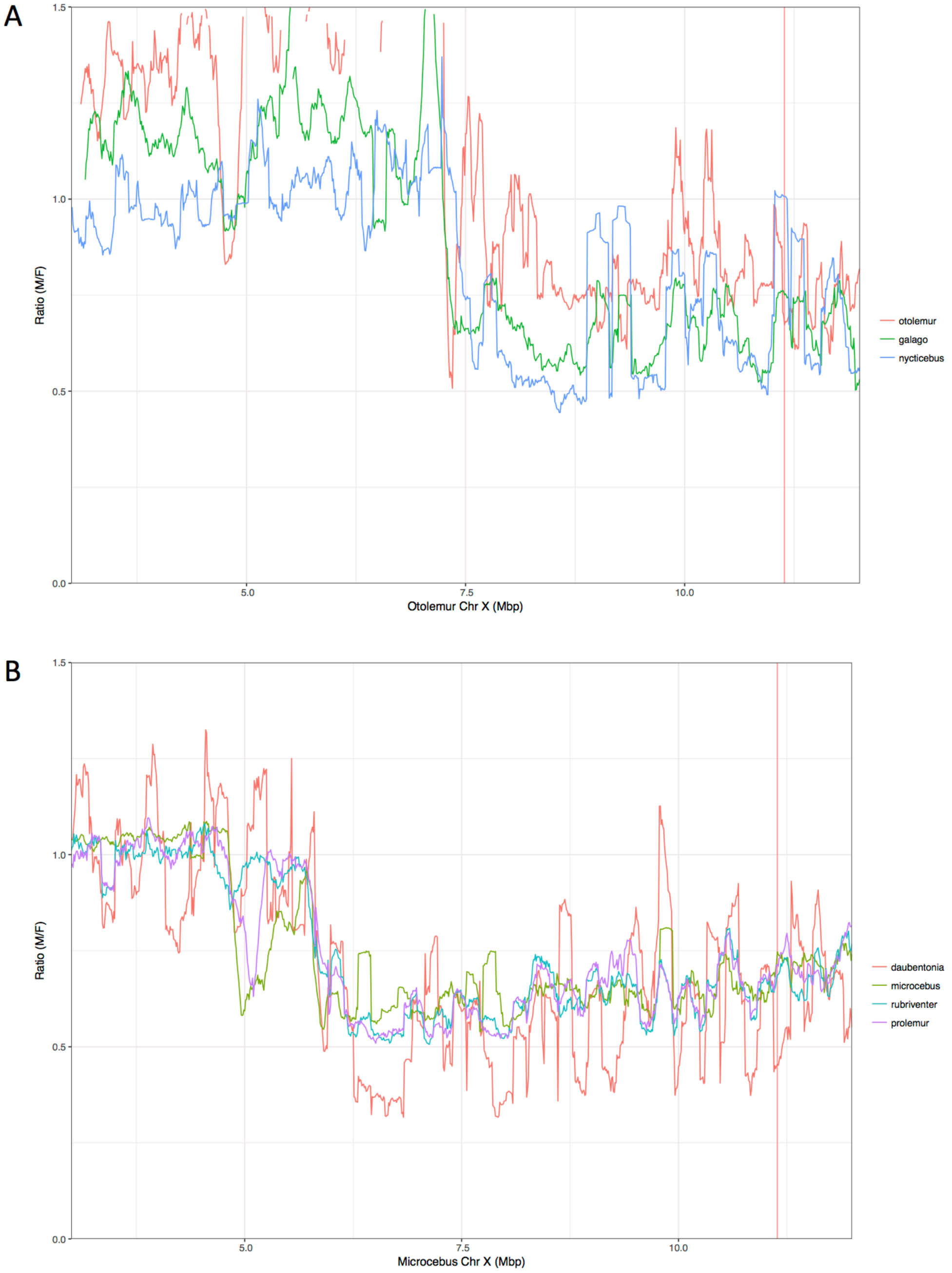

**Fig. S3.**

SNP density analysis. M:F SNP density ratio (ln scale) for all seven strepsirrhine species (see Methods for details). Dashed lines indicate the mean m:f SNP density across sliding windows of the same size on chromosome 4, the 97.5% and 2.5% quantiles, to show the variation across the autosomes. See legend of figure 1 for more details.

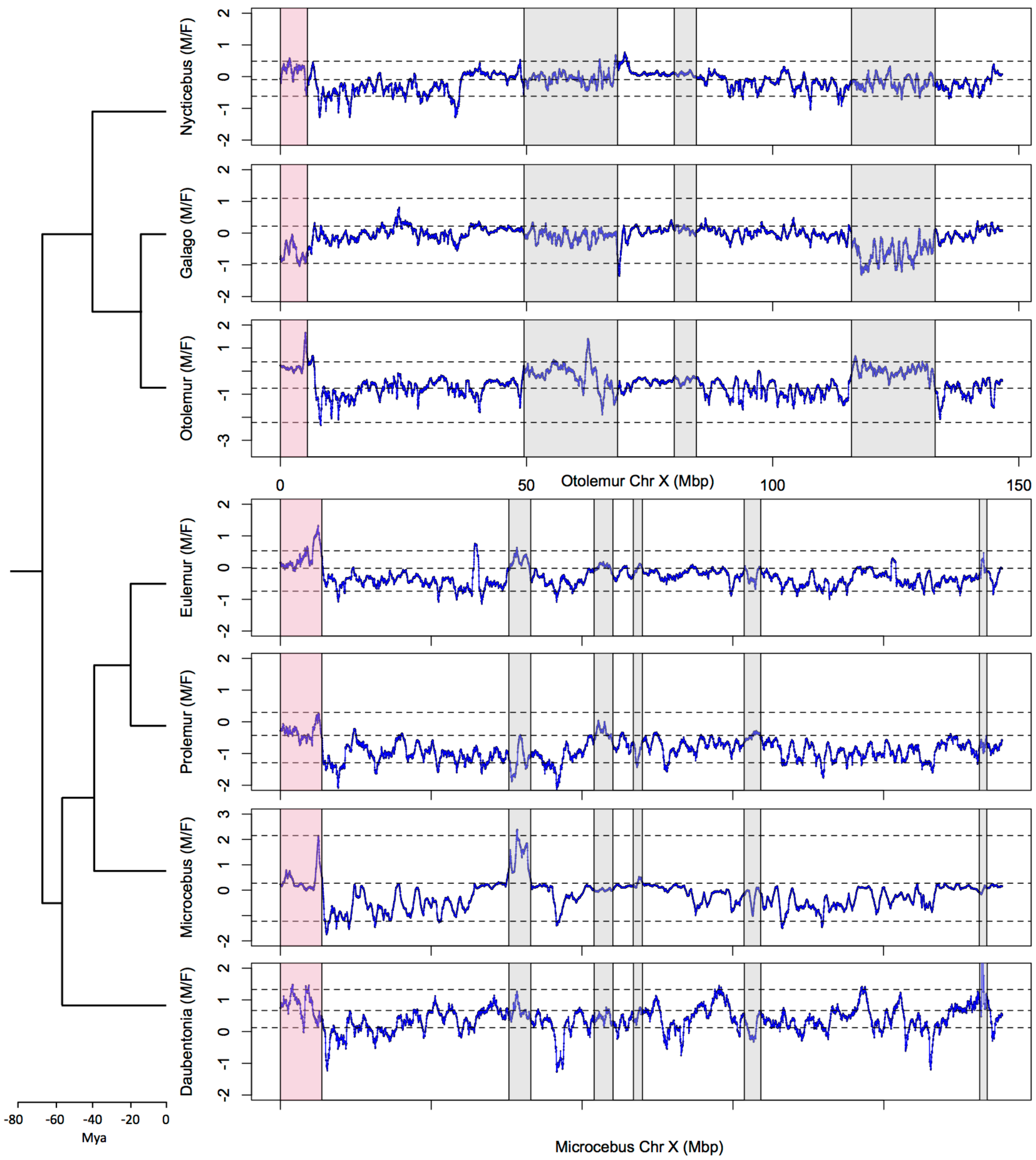

**Table S1.**
Statistics about the genome sequencing in the 7 species.

| Lineage | Genus | Species | Common name | Sex | Read # | Read length | Sequences (Gb) | Coverage (X)* | Source |
|---|---|---|---|---|---|---|---|---|---|
| Lemuriformes | *Daubentonia* | *madagascariensis* | aye-aye | M | 574 860 296 | 125 | 71.9 | 23.2 | MNHN, Paris |
| Lemuriformes | *Daubentonia* | *madagascariensis* | aye-aye | F | 807 533 380 | 150 | 121.1 | 39.1 | Zoo Frankfurt |
| Lemuriformes | *Microcebus* | *murinus* | gray mouse lemur | M | 553 217 340 | 125 | 69.2 | 22.3 | MNHN, Brunoy |
| Lemuriformes | *Microcebus* | *murinus* | gray mouse lemur | F | 567 375 076 | 125 | 70.9 | 22.9 | MNHN, Brunoy |
| Lemuriformes | *Eulemur* | *rubriventer* | red-bellied lemur | M | 361 251 832 | 150 | 54.2 | 17.5 | Zoo de Lyon |
| Lemuriformes | *Eulemur* | *rubriventer* | red-bellied lemur | F | 316 639 574 | 150 | 47.5 | 15.3 | Zoo de Lyon |
| Lemuriformes | *Prolemur* | *simus* | greater bamboo lemur | M | 242 884 578 | 150 | 36.4 | 11.8 | Zoo de Lyon |
| Lemuriformes | *Prolemur* | *simus* | greater bamboo lemur | F | 428 087 286 | 150 | 64.2 | 20.7 | Zoo de Lyon |
| Lorisiformes | *Nyctibebus* | *coucang* | slow loris | M | 665 798 842 | 150 | 99.9 | 32.2 | MNHN, Paris |
| Lorisiformes | *Nyctibebus* | *coucang* | slow loris | F | 670 569 564 | 150 | 100.6 | 32.4 | MNHN, Paris |
| Lorisiformes | *Galago* | *senegalensis* | senegal bushbaby | M | 641 724 580 | 150 | 96.3 | 31.1 | MNHN, Paris |
| Lorisiformes | *Galago* | *senegalensis* | senegal bushbaby | F | 666 087 196 | 150 | 99.9 | 32.2 | MNHN, Paris |
| Lorisiformes | *Otolemur* | *garnetti* | northern greater galago | M | 662 474 900 | 150 | 99.4 | 32.1 | MNHN, Paris |
| Lorisiformes | *Otolemur* | *garnetti* | northern greater galago | F | 2 599 993 104 | 100 | 260.0 | 83.9 | EBI** |

*based on human genome size (assuming similar genome sizes in humans and all these species)
**SRR016877 to SRR016896 files fastq.gz 1 and 2.
